# Single nucleus transcriptomic analysis of human dorsal root ganglion neurons

**DOI:** 10.1101/2021.07.02.450845

**Authors:** Minh Q. Nguyen, Lars J. von Buchholtz, Ashlie N. Reker, Nicholas J. P. Ryba, Steve Davidson

## Abstract

Somatosensory neurons with cell bodies in the dorsal root ganglia (DRG) project to the skin, muscles, bones, and viscera to detect touch and temperature as well as to mediate proprioception and many types of interoception. In addition, the somatosensory system conveys the clinically relevant noxious sensations of pain and itch. Here we used single nuclear transcriptomics to characterize the classes of human DRG neurons that detect these diverse types of stimuli. Notably, multiple types of human DRG neurons have transcriptomic features that resemble their mouse counterparts although expression of genes considered important for sensory function often differed between species. More unexpectedly, we demonstrated that several classes of mouse neurons have no direct equivalents in humans and human specific cell-types were also identified. This dataset should serve as a valuable resource for the community, for example as means of focusing translational efforts on molecules with conserved expression across species.

## Introduction

The somatosensory system responds to a wide range of mechanical, thermal and chemical stimuli to provide animals with critical information about their environment and internal state. For example, our sense of touch is mediated by mechanosensory neurons with somata located in the dorsal root and trigeminal ganglia that innervate the skin (*1*). In addition to the skin, somatosensory neurons target specialized sensory environments like the cornea and conjunctiva or meninges (*2, 3*), the internal organs (*4*) as well as bones and muscles to provide rich perceptual experiences and trigger appropriate behavioral, reflex and autonomic responses (*5*). Amongst their many roles, somatosensory neurons provide input for the conscious perception of pain and itch (*6, 7*) and the subconscious coordination of muscles and limbs known as proprioception (*8*). Peripheral neurites of somatosensory receptor cells must adapt to growth, reinnervate targets after injury and are also affected by inflammation (*9*).

Studies in model organisms have characterized a range of sensory and growth factor receptors and ion-channels that contribute to the properties and selectivity of somatosensory neurons (*10–12*). Some of these, like the cooling and menthol sensing receptor (*Trpm8*) appear to define functional classes of cells (*13*). By contrast, the sense of touch appears to use a complex distributed code involving several different types of cells (*14*) to achieve its remarkable discriminatory power. For the most part, the human somatosensory system expresses the same range of functional genes as rodents (*15*) and exhibits similar responses to many types of stimulus (*6, 8, 10, 12, 16, 17*). Moreover, rare individuals with loss of function variants of several of these genes have deficits that recapitulate key effects of knocking out that gene in mice (*8, 18–21*). However, despite the identified similarities between mice and humans, the success of translating new therapeutic strategies that are effective for treating pain in mice has often been disappointing when tested in human subjects (*22, 23*).

Recently, various directed genetic strategies have been used in mice to characterize the response properties and anatomical features of a variety of interesting classes of large diameter, fast conducting Aβ- and Aδ-subtypes (*1*). Interestingly, these neurons generally have complex peripheral endings that often target hair follicles. Human skin hairs are quite different from those in mice, suggesting that there may be significant differences between the large diameter neurons in mice and humans. By contrast, most types of small diameter, slow conducting c-fibers terminate as free nerve endings both in mice and humans (*6*). Single cell sequencing approaches have produced a transcriptomic classification for mouse somatosensory neurons that corresponds well with their anatomy and function (*5, 14, 24, 25*). Intriguingly, in mice members of the Mrgpr-family of GPCRs mark at least two classes of small diameter neurons (*24, 25*). Mrgprs have undergone massive genetic expansion in rodents, not seen in other animals, often making it difficult to identify true orthologs in humans (*26, 27*). A map of human somatosensory neuron transcriptomic classes would help uncover selective differences between the sensory neurons in mice and humans and provide clues as to how similar somatosensory input is in the two species. Finally, such analysis may provide important new targets to consider for translational approaches to treat both pain and itch. Here we used nuclei based single cell transcriptomics to generate a comprehensive description of human cell types, highlight similarities and surprising differences between somatosensory neuron classes in humans and mice that are reflected not only in terms of individual genes but can be discerned in co-clustering. We used multigene *in situ* hybridization (ISH) to help confirm these conclusions and present evidence for anatomic organization of functionally distinct neuronal classes in the human dorsal root ganglion.

## Results

### Generating a representative transcriptomic map of human somatosensory cell types

Single lumbar L4 and L5 human dorsal root ganglia were rapidly recovered from transplant donors within 90 minutes of cross-clamp and were immediately stored in RNAlater. Nuclei from individual ganglia were isolated and samples were enriched for neuronal nuclei by selection using an antibody to NeuN. Five ganglia from one male and four female donors with ages ranging from 34 to 55 were subjected to droplet based single nucleus (sn) capture, barcoding, and reverse transcription (10X Genomics). Combinatorial clustering methods (*28*) allowed co-clustering of neuronal nuclei into well-defined and distinct transcriptomic groups from their non-neuronal counterparts (Figure 1A, Figure 1-figure supplement 1). After removal of non-neuronal nuclei from the dataset, re-clustering the DRG-neuron data identified a range of more than a dozen diverse transcriptomic classes of human somatosensory neurons (Figure 1B, Figure1-figure supplement 1).

**Figure 1.**
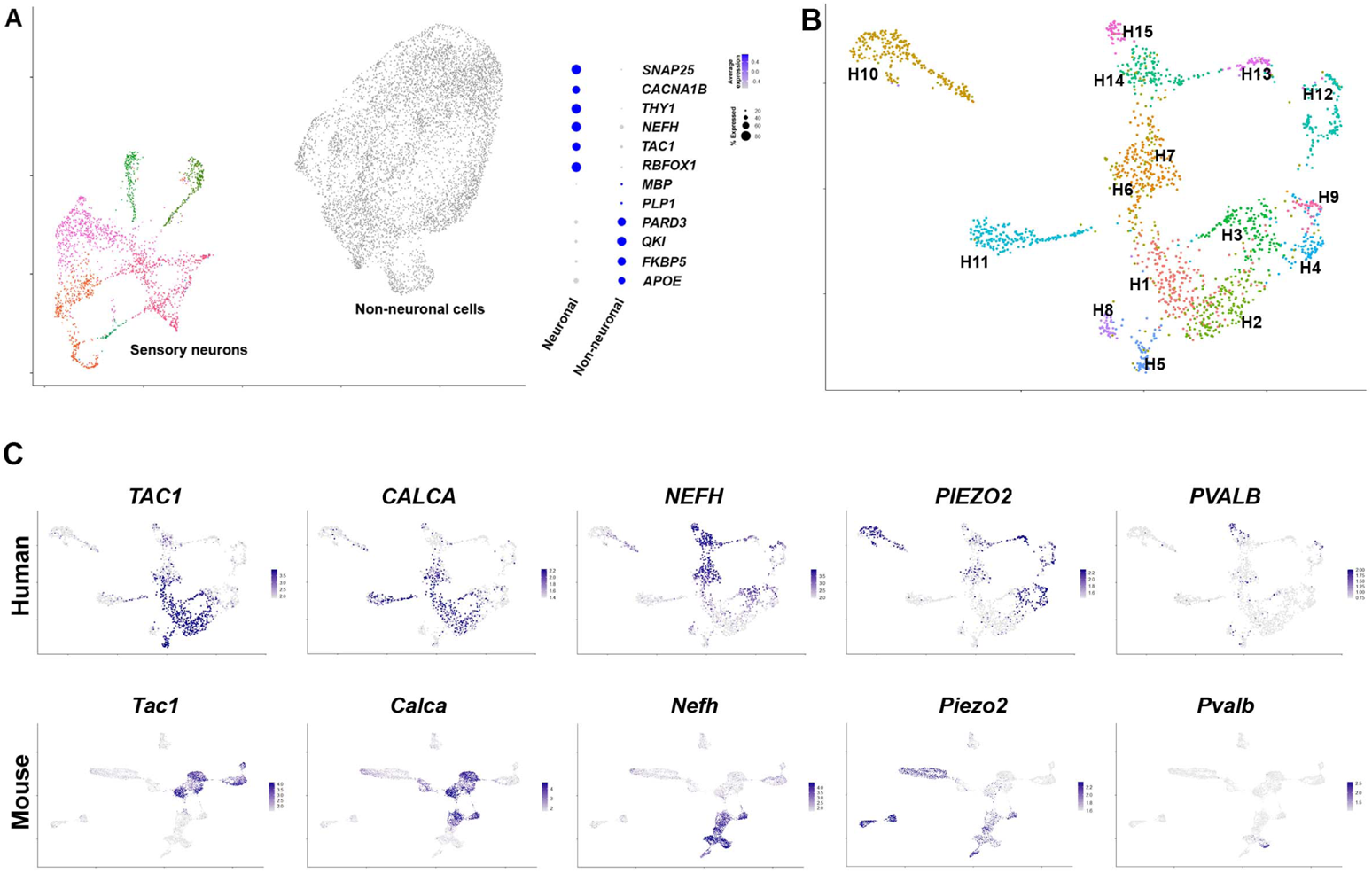
Diverse classes of human DRG neurons revealed by single nuclear transcriptomics. (**A**) Universal manifold (UMAP) representation of graph-based co-clustered snRNA-sequences from human DRG nuclei reveal two well separated groups corresponding to sensory neurons (colored) and non-neuronal cells (gray). To the right, a dot-plot highlights the expression of markers that can be used to distinguish these groups of cells (see also Figure 1-figure supplement 1A). (**B**) Reanalyzing 1837 neuronal nuclei identifies fifteen types of human DRG neurons that have been differentially colored. Transcriptomic similarity to mouse neuronal types allows tentative classification of some of these neuronal classes. (**C**) UMAP representation of human DRG neurons showing relative expression level (blue) of diagnostic markers. For comparison UMAP representation of mouse neurons (*29*) showing the relative expression patterns of the same markers (see Figure 1-figure supplement 1 for more details). In combination, the expression patterns of these genes and other markers (Figure 1-figure supplements 1–3) were used to tentatively classify cell classes.

One of the best studied groups of somatosensory receptors in mice are nociceptive peptidergic neurons that co-express a variety of neuropeptides including substance P, calcitonin gene related peptide (CGRP) and pituitary adenylate-cyclase-activating polypeptide (PACAP). These neurons are typically small soma diameter, non-myelinated, slow conducting c-fibers, but also include faster conducting lightly myelinated Aδ-neurons (*24, 25*). In the human DRG dataset, *TAC1* (substance P), *CALCA* and *CALCB* (CGRP) and *ADCYAP1* (PACAP), are expressed in several transcriptomic classes (H1, H2, H3, H5, H6, Figure 1C, Figure 1-figure supplements 2, 3). For comparison the expression of the same genes in mouse DRG neurons is shown (Figure 1C, Figure 1-figure supplements 2, 3) using data from single nuclei sequencing (*29*). Just as in mice, the putative human peptidergic nociceptors express the high affinity nerve growth factor receptor *NTRK1*, the capsaicin and mustard oil gated ion channels *TRPV1* and *TRPA1* but generally only low levels of the stretch gated ion channel *PIEZO2* (Figure 1C, Figure 1-figure supplements 2, 3).

Although previous localization studies have suggested that in humans the neurofilament protein *NEFH* is expressed in all sensory neurons (*30*), this gene showed graded expression in our data (Figure 1C) and marks several classes of cells just as in mice (Figure 1-figure supplement 3). Some of these (including H3 and H6) also express peptidergic markers and the pain related voltage gated sodium channel *SCN10A* (Figure 1-figure supplement 3) and thus have molecular hallmarks of Aδ-nociceptors (*2*). However, the neuronal classes H14 and H15 expressing the highest levels of *NEFH* are distinct from the peptidergic neurons (Figure 1C, Figure 1-figure supplements 2, 3), likely representing different types of large diameter, fast conducting myelinated Aβ-neurons. These cell types are neurotrophin 3 receptor *NTRK3* positive, some also contain the brain derived neurotrophic factor receptor *NTRK2* but exhibit little expression of *NTRK1* (Figure 1-figure supplements 2, 3). In mice, proprioceptors are a subtype of Aβ-neurons marked by the calcium binding protein parvalbumin, the transcription factor *Etv1* and the voltage gated sodium channel subunits *Scn1a* and *Scn1b* (*24, 29*). In the human data, the small H15 group of *NTRK3*-positive cells had this expression pattern (Figure 1C, Figure 1-figure supplement 3) implying that proprioceptors have conserved transcriptomic markers in humans and mice. Similarly, small groups of both Aδ-low threshold mechanosensors (H13) and cool responsive neurons (H8) were identified by their characteristic expression profiles of functionally important transcripts (Figure 1-figure supplement 3). Thus, large groups of human and mouse DRG neurons appear to share basic transcriptomic signatures and functional potential, supporting our data as informative about neuronal diversity amongst human somatosensory neurons.

Despite these similarities between the putative peptidergic, proprioceptive, cooling sensitive, Aβ- and Aδ-classes of DRG neurons in mice and humans there were important differences in their expression of genes that may be functionally significant. These include molecules that modulate cellular responses to internal signals (e.g. growth factor receptors), sensory stimuli and also the mediators they may release. For example, in humans, the H8 putative cool responsive neurons expressing *TRPM8* were strongly positive for the BDNF-receptor *NTRK2* but hardly expressed the neuropeptide *TAC1* whereas in rodents the converse was true (Figure 1-figure supplements 2, 3). Other genes that have been shown to control sensory responses in mice exhibit a different expression pattern in human DRG neurons. For instance, *Tmem100* encodes a protein that in mice has been implicated as playing an important role in functional interactions between *Trpv1* and *Trpa1* and contributing to persistent pain (*31*). By contrast it was almost undetectable in the human sequencing data (Figure 2A). Similarly, we did not detect marked expression of the sphingosine-1-phosphate receptor *S1PR3* (Figure 2A) that has been suggested as a target for treating both pain and itch based on mouse work (*32*). More strikingly, a small group of human neurons, H5, expressing *TRPA1* were resolved in our clustering (Figure 2B), whereas in mouse nuclear sequencing data no direct counterpart was detected (Figure 1-figure supplement 2). Interestingly cell-based sequencing (*24*) of mouse DRG neurons does identify a group of peptidergic nociceptors (called CGRP-gamma) with abundant *Trpa1* expression, highlighting the risk of over-interpreting differences across species.

**Figure 2.**
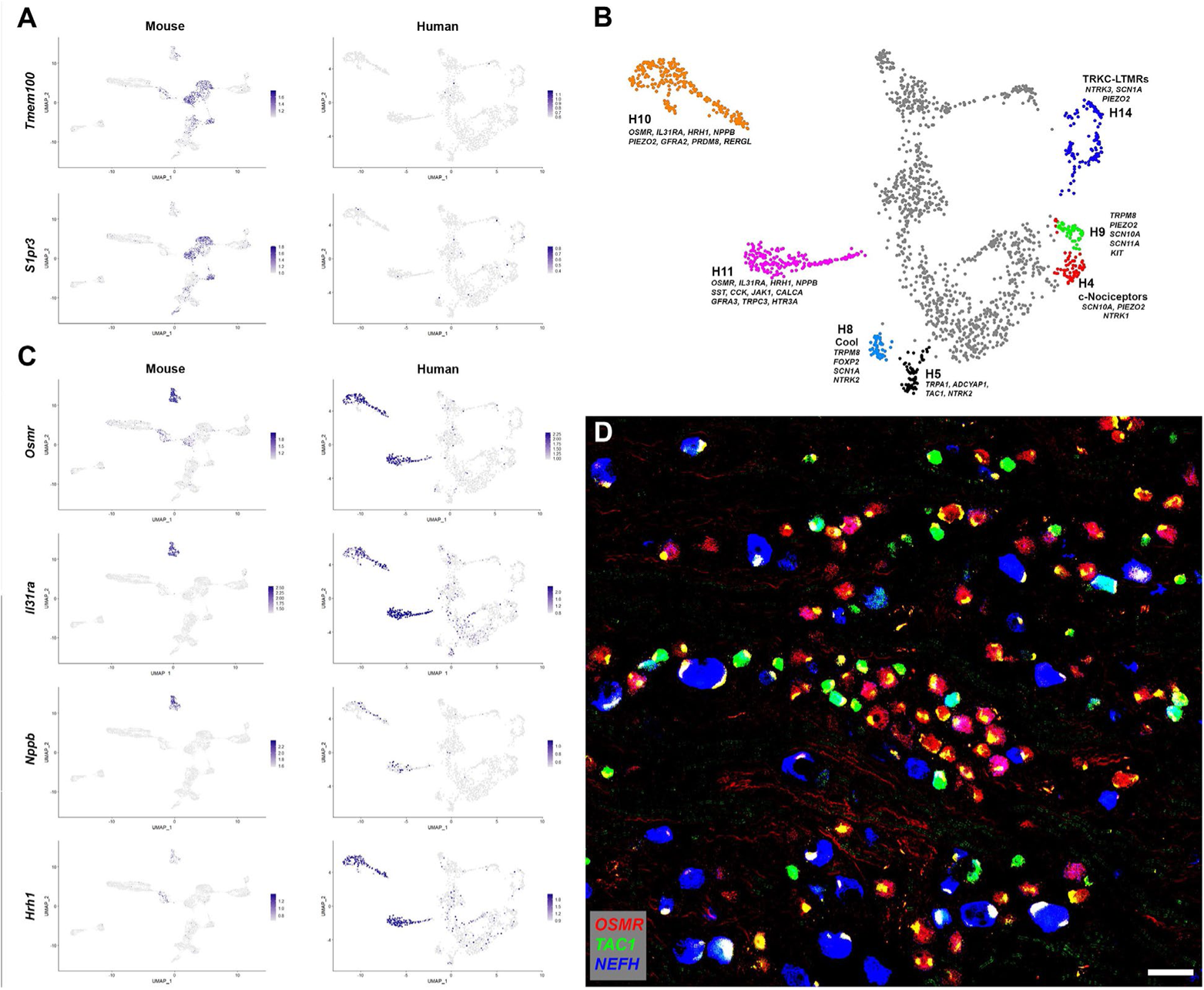
Human DRG neurons exhibit specialization that distinguishes them from mouse counterparts. (**A**) UMAP representation of mouse and human DRG neurons showing relative expression level (blue) of two genes that have been linked to pain sensation in mice. Note that both *TMEM100* and *S1PR3* are more sporadically expressed by the human somatosensory neurons and are not markers of select cell types. (**B**) Classes of DRG neurons that are selectively detected in humans are highlighted together with their expression of key genes. H9 neurons co-express the cool and mechanosensory ion channels; for comparison cool sensitive neurons (H8) that correspond more closely with their rodent counterparts are also highlighted. (**C**) Expression profiles of select itch related genes in the mouse and human DRG transcriptome. (**D**) Confocal image of a region from a human DRG that was labeled using multiplexed ISH for *OSMR*, *TAC1* and *NEFH* as indicated in the key. Almost all neurons detected by any of the 9 ISH probes (see Methods) were *OSMR*, *TAC1* or *NEFH* positive: only 62 out of 1153 cells (5.4 %) in 2 complete sections were not positive for one of these three genes. However, few neurons were strongly positive for more than one of these markers (see Figure 2-figure supplement 1 for individual channels). Note that autofluorescence in all channels from lipofuscin associated with many human neurons appears white in the overlay image and should not be confused with real signal (see Figure 2-figure supplement 1 for more detail). Also note that *NEFH* is typically expressed in larger diameter neurons than the other two markers. Scale bar = 100 µm.

Nonetheless, whereas mouse CGRP-gamma neurons strongly express *Calca* and *Ntrk1*, H5 cells are essentially *CALCA* (CGRP) and *NTRK1* negative and instead are strongly *NTRK2* positive (Figure 1, Figure 1-figure supplements 2, 3) suggesting that they may respond differently to external stimuli and in their signaling properties. Thus, the availability of human transcriptomic data should help focus translational work in model organisms on promising targets with conserved expression patterns in humans.

### Human DRG neurons without clear transcriptomic equivalents in mice

Analysis of the gene expression patterns of the different classes of human somatosensory neurons revealed several groups for which we could not discern direct counterparts in the mouse. One small but prominent group of human DRG neurons (H9) expresses *TRPM8*, *PIEZO2*, *SCN10A* and *SCN11A* (Figure 1-figure supplements 2, 3, Figure 2B) and clearly segregates from the putative cool sensing cells (H8) that express *TRPM8, GPR26, NTM* and *FOXP2* but are devoid of both the light touch receptor and the pain related sodium channels (Figure 2B, Figure 1-figure supplement 3). In mice, *Trpm8* expression is simpler with the cool sensing, menthol responsive ion channel just expressed in cells with this latter gene expression pattern (Figure 1-figure supplement 3). Interesting single fiber recordings have identified human neurons that respond to both cooling and gentle touch as might be expected for cells expressing both *TRPM8* and *PIEZO2* (*33*). H9 neurons resemble (but also have differences from) human mechanosensory neurons that were recently engineered by transcriptional programming of stem cells (*34*).

A second larger group of human neurons H12 is marked by *NTRK3* and the voltage gated ion channel *SCN1A*, but is only weakly positive for *NEFH*, expresses moderate levels of *PIEZO2* (Figure 2B, Figure 1-figure supplement 3) and appears distinct from any potential mouse counterpart. The H12 gene expression pattern is most consistent with these cells functioning as a type of mechanosensor that has no direct equivalent in mice. Similarly, we designated H4 as c-nociceptors because of their expression of nociception related *SCN10A* and *NTRK1* and low level of *NEFH* (Figure 2B, Figure 1-figure supplement 3). These neurons expressed low levels of neuropeptides, but their overall gene expression patterns did not resemble any mouse counterparts including the non-peptidergic nociceptors (see below).

The two remaining large groups of neurons in the human dataset H10 and H11 that have no clear mouse counterpart exhibit most similarity with mouse c-type non-peptidergic neurons (Figure 2-figure supplement 1A). At a functional level both H10 and H11 express receptors that in mice have roles in detecting pruritogens. For example, these clusters were positive for the two subunits (*IL31RA* and *OSMR*) of the interleukin 31 receptor and the histamine receptor *HRH1* (Figure 2C) that mediate mast cell related scratching in mice (*35*). They also express the itch related neuropeptide *NPPB* (Figure 2C), nociception related sodium channels *SCN10A* and *SCN11A* as well as *TRPV1* (Figure 2-figure supplement 1A) but not appreciable *NEFH* or *TAC1* (Figure 1C). Therefore, it is likely that these are groups of putative unmyelinated, non peptidergic nociceptors with roles in triggering human itch responses.

The peptidergic nociceptors, myelinated Aβ and Aδ-neurons, rarer human specific cells, and the two non-peptidergic nociceptor clusters H10 and H11 account for all the neurons in our analysis with H10 and H11 totaling approx. 20% of the neurons. In marked contrast, mouse non-peptidergic, small diameter neurons are far more numerous than H10 and H11 accounting for 40% or the sensory neurons in mouse DRGs (*29*) and divide into 4 highly stereotyped transcriptional groups (Figure 1-figure supplement 2). Two of these classes of mouse neurons (NP2 and NP3) trigger itch (*7, 36*), one (NP1, expressing *Mrgprd*) responds to noxious mechanical stimulation (*14*). NP1 neurons may have a role in mechano-nociception (*5*) and have recently been associated with suppression of skin inflammation (*37*), which was hypothesized as relevant for human health. The fourth class corresponds with low threshold mechanosensors (cLTMRs) that are thought to mediate affective touch (*5, 38*). Given this difference between the transcriptomic map of human DRG neurons and their rodent counterparts, we next used independent ISH-based analysis to test basic predictions of the sequencing. If transcriptomic characterization of human DRG neurons is accurate then one clear expectation is that *TAC1*, *NEFH* and *OSMR* should be expressed by distinct and only partially overlapping populations of human DRG neurons. If it is also comprehensive, then we would anticipate that the same three markers should label the vast majority of neurons. Multigene ISH demonstrates that both these predictions are true for human DRG neurons (Figure 2D, Figure 2-figure supplement 1B) with essentially every cell labeled by one of these probes but with very few exhibiting strong co-expression. Although *NEFH* expression could be detected in some of the cells positive for the other markers (Figure 2D), many *TAC1* or *OSMR*-positive small diameter neurons were negative for this neurofilament subunit. Moreover, *TAC1* and *OSMR* labeled almost completely separate sets of cells. Notably, in keeping with our assignments based on transcriptomic data, the largest diameter neurons are strongly positive for *NEFH* whereas *TAC1* and *OSMR* primarily label smaller cells (Figure 2D). Finally, in keeping with snRNA-sequence analysis, these three markers each labeled a large group of neurons.

### Co-clustering human and mouse DRG neuron snRNAseq data

As detailed above, the expression of genes that are important for functional and morphological features of somatosensory neurons reveal similarities between groups of human and mouse neurons. They also expose differences that likely reflect distinct somatosensory adaptations in the two species. We next used co-clustering methods to test whether the wider transcriptome could reveal additional information about the relationships between classes of human and rodent DRG neurons using the same mouse dataset (*29*) that we analyzed above (Figure 1, Figure 1-figure supplement 2). We used the well-established approach developed by the Satija lab (*28*) as it has been shown to perform well without forcing false class assignments. As predicted, several classes of human neurons grouped with corresponding mouse counterparts including H15 – proprioceptors, H14 – Aβ cells, H13 – Aδ-LTMRs, H11 – NP3 (*Nppb*) neurons and H3/H6 – Aδ-nociceptors (Figure 3A). This analysis suggested that H10 the other cluster that gene expression indicated are also itch related most closely resembled NP1 (*Mrgprd*) neurons rather than any other human or mouse class of sensory neurons. The H12 cluster, which is human specific, grouped close to larger diameter mouse neurons, whereas other clusters of human cells appeared better aligned with smaller diameter nociceptors. However, all types of peptidergic small diameter nociceptors were less organized in the co-clustering and separated from their potential mouse counterparts despite their qualitatively similar expression of functional markers (Figure 1-figure supplement 2).

**Figure 3.**
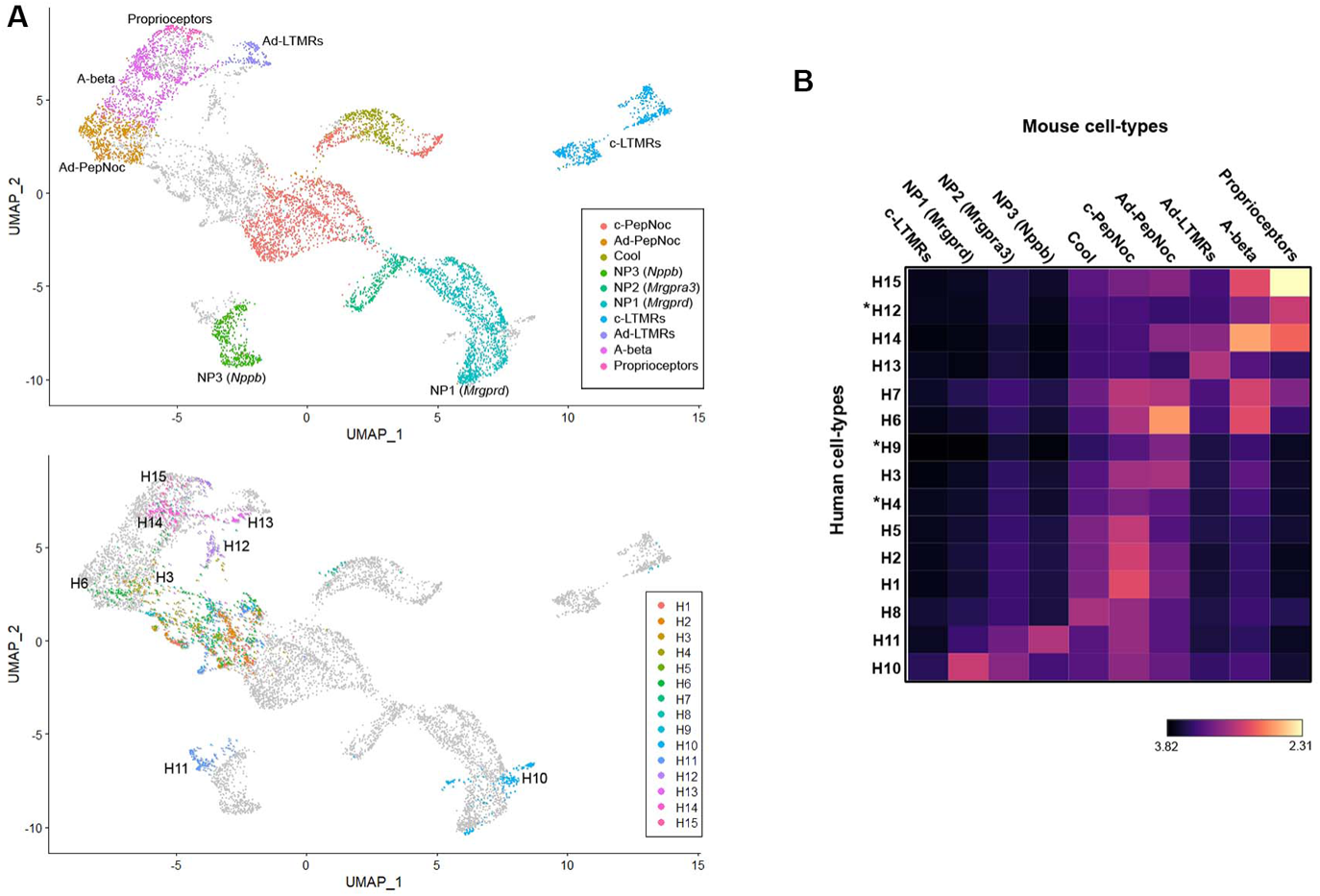
Co-clustering of human and mouse neurons largely tentative assignments based on select genes. (**A**) UMAP representation of the co-clustering of mouse and human neurons. Upper panel shows the mouse neurons colored by their identity when analyzed alone (Figure 1-figure supplement 1); lower panel shows human neurons colored by their identity when analyzed alone (Figure 1). Note that large diameter human neurons match their expected mouse counterparts reasonably well and that the two classes of neurons expressing itch related transcripts H10 and H11 best match NP1 and NP3 neurons, respectively. (**B**) Heatmap showing the natural logarithm (see scalebar) of Kullback-Leibler divergences for the various human neuron classes when compared to each class of mouse cells as a reference distribution; human specific classes are marked by *.

UMAP plots (Figure 3A) provide a visual representation of similarity between cells with related transcriptomic properties. However, since they collapse multidimensional information into two dimensions, relationships between separated clusters are harder to interpret. Therefore, we made use of Kullback-Leibler divergence estimation to quantitate the similarity between human DRG neuron clusters and all their potential mouse counterparts (Figure 3B). As expected, clusters that co-segregate in the UMAP analysis showed greatest similarity but additional relationships not apparent from the visual representation of the co-clustering were also seen. For example, the small cluster of human “cool” responsive neurons H8 showed greatest similarity to mouse Trpm8-cells and several groups of human cells (H1, H2, H5) that gene expression predicted should be c-type peptidergic nociceptors, indeed best matched these cells (Figure 3B). Interestingly, no class of human neurons showed appreciable similarity to mouse c-LTMRs.

Amongst the groups of cells that had human specific gene expression patterns, H9 (the putative cool and mechanical responsive cells) showed only weak similarity to any mouse neuron class. H12, which we considered likely to be mechanosensors best matched mouse proprioceptors and H4 neurons appeared distantly related to several classes of nociceptor but without a clear match in mice. One important caveat to this type of analysis remains that any functional conclusions based on shared transcriptomic features still need to be verified experimentally.

### Transcriptomically related neurons are spatially grouped in the human dorsal root ganglion

From sequence analysis we identified a range of potential markers to better explore the diversity of human DRG neurons using ISH. To maximize information, we chose a highly multiplexed approach (Figure 4) revealing the different classes of sensory neurons identified in the transcriptomic data. For example, *TRPM8* expressing neurons clearly segregate into two distinct types (Figure 4A, Figure 4-figure supplement 1). One set of cells (H8) share other transcriptomic properties with mouse cooling responsive cells. For example, in H8 neurons, *TRPM8*, the cool and menthol receptor is not co-expressed with the ion channels *SCN10A* or *PIEZO2* (Figure 4A) but these cells are *NTRK2* positive (Figure 4-figure supplement 1). By contrast, other cells (H9) co-express the pain and light touch related ion channels (*SCN10A* and *PIEZO*) with *TRPM8* (Figure 4A, Figure 4-figure supplement 1). Similarly, putative proprioceptive neurons (H15) were distinguished by their expression of *NEFH*, *PIEZO2* and *PVALB* and lack of *NTRK2* (Figure 4B, Figure 4-figure supplement 1). One surprise (Figure 4A, B) was that in small fields of view, several examples of all three of these rare neuron types could be identified in human DRGs. By contrast, much of the rest of the ganglion was devoid of these cell types and instead the neurons there had distinct sets of markers. Therefore, it appears that transcriptomic classes of human DRG sensory neurons may not be stochastically distributed in the ganglion as is thought to be the case in mice. Indeed, when we examined the distribution of nociceptors and myelinated neurons at lower magnification (using strong selective probes), broad clustering of similar types of neurons was apparent, quantifiable, and statistically significant (Figure 4C, Figure 4-figure supplement 1).

**Figure 4.**
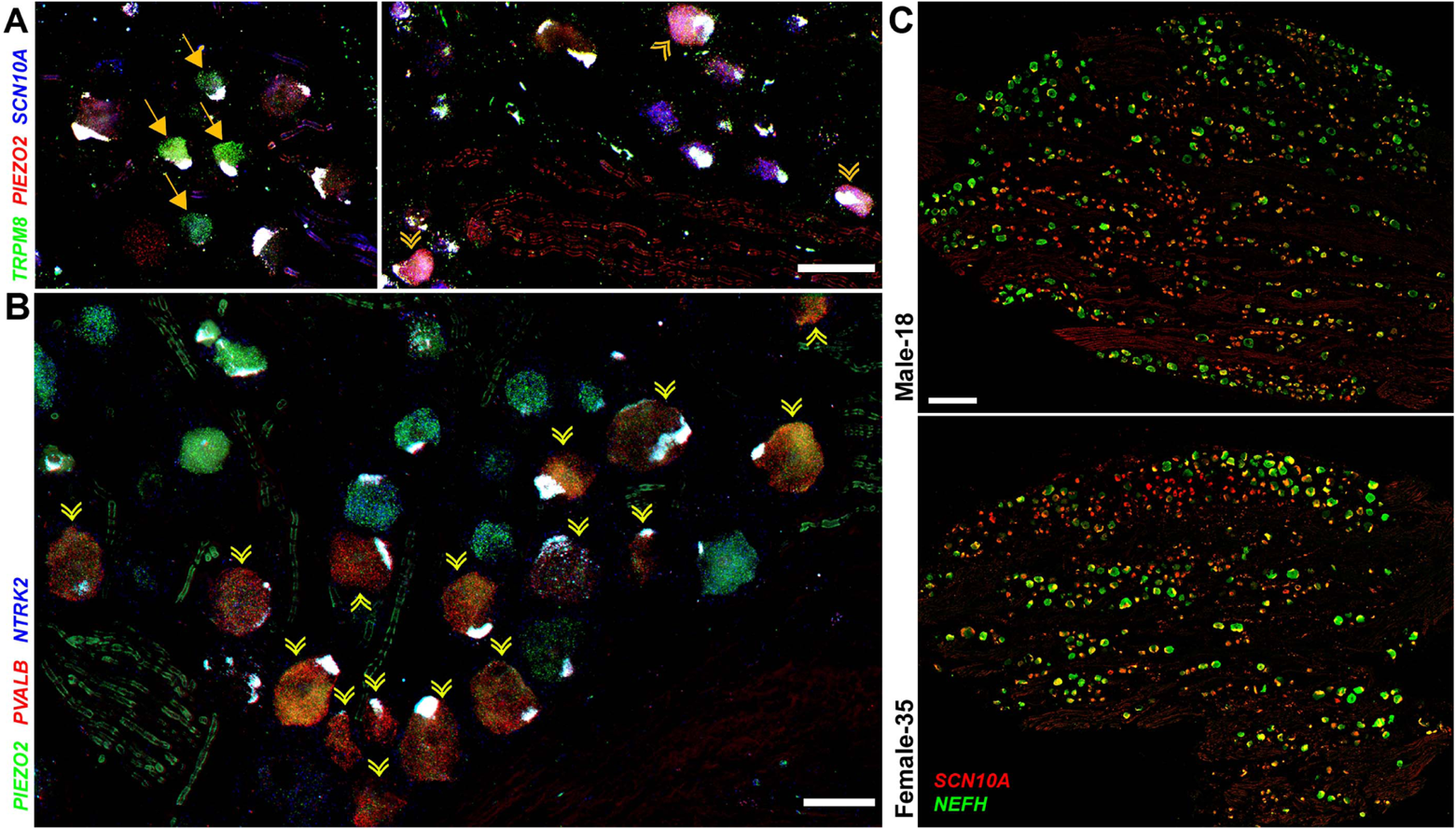
Transcriptomically related classes of human DRG neurons are spatially clustered in the ganglion. Confocal images of sections through a human DRG probed for expression of key markers using multiplexed ISH; see Figure 4-figure supplement 1 for the individual panels and additional probes. (**A**) Left panel shows a group of four cool neurons (yellow arrows) that express *TRPM8* (green) but not *PIEZO2* (red) or *SCN10A* (blue). By contrast, right panel shows a different region of the ganglion where three CM neurons co-express these three transcripts (double arrowheads). (**B**) Other regions of the ganglia were dominated by larger diameter neurons. Putative proprioceptors, highlighted by double arrowheads, expressing *PIEZO2* (green) and *PVALB* (red), but not *NTRK2* (blue) were typically highly clustered in the ganglion. (**C**) Lower magnification images of complete sections stained for *NEFH* (green) and *SCN10A* (red) highlight the extensive co-clustering of large and small diameter neurons in different individuals (see Figure 4-figure supplement 1 for quantitation). Scale bars = 100 µm in (**A**) and (**B**); 500 µm in (**C**).

### H10 and H11 are distinct but related types of human nonpeptidergic neurons

Perhaps the most intriguing classes of human somatosensory neurons revealed by our transcriptomic approach are the H10 and H11 classes that primarily share features with the mouse non-peptidergic nociceptors NP1-3 (Figures 2, 3, Figure 2-figure supplement 1, Figure 5-figure supplement 1). ISH showed that the H10 and H11 classes of neurons, identified by their expression of *OSMR* were small diameter neurons comparable in size to the *TAC1*-expressing peptidergic nociceptors (Figure 5A). Interestingly two qualitatively different ratios of *SCN10A* and *OSMR* were apparent in these cells (Figure 5A) hinting at their distinct identities. Our data (Figure 2, Figure 2-figure supplement 1) show that H10 and H11 neurons express a number of genes that are known to be expressed in mouse NP3 cells and functionally important for triggering pruritic responses (*5, 35*). They are also distinguished from each other by expression of genes that likely play roles in itch and other aspects of somatosensation (Figure 5B, Figure 2-figure supplement 1, Figure 5-figure supplement 1). For example, although not prominently expressed, the human chloroquine responsive receptor *MRGPRX1* (*27*) localized selectively to H10 neurons (Figure 5B) perhaps suggesting a relationship to mouse NP2 cells. By contrast, Janus kinase 1 (*JAK1*), a mediator of itch through various types of cytokine signaling (*39*), including through OSMR, and the neuropeptide *SST* are particularly strongly expressed in H11 cells (Figure 5B). Both these genes are prominent markers of NP3 pruriceptors in mouse (Figure 5B). However, not all known itch related transcripts are expressed in H10 and H11 neurons and both classes of cells express genes that better define NP1 neurons in mice as well as other cell types (Figure 5B, Figure 2-figure supplement 1, Figure 5-figure supplement 1).

**Figure 5.**
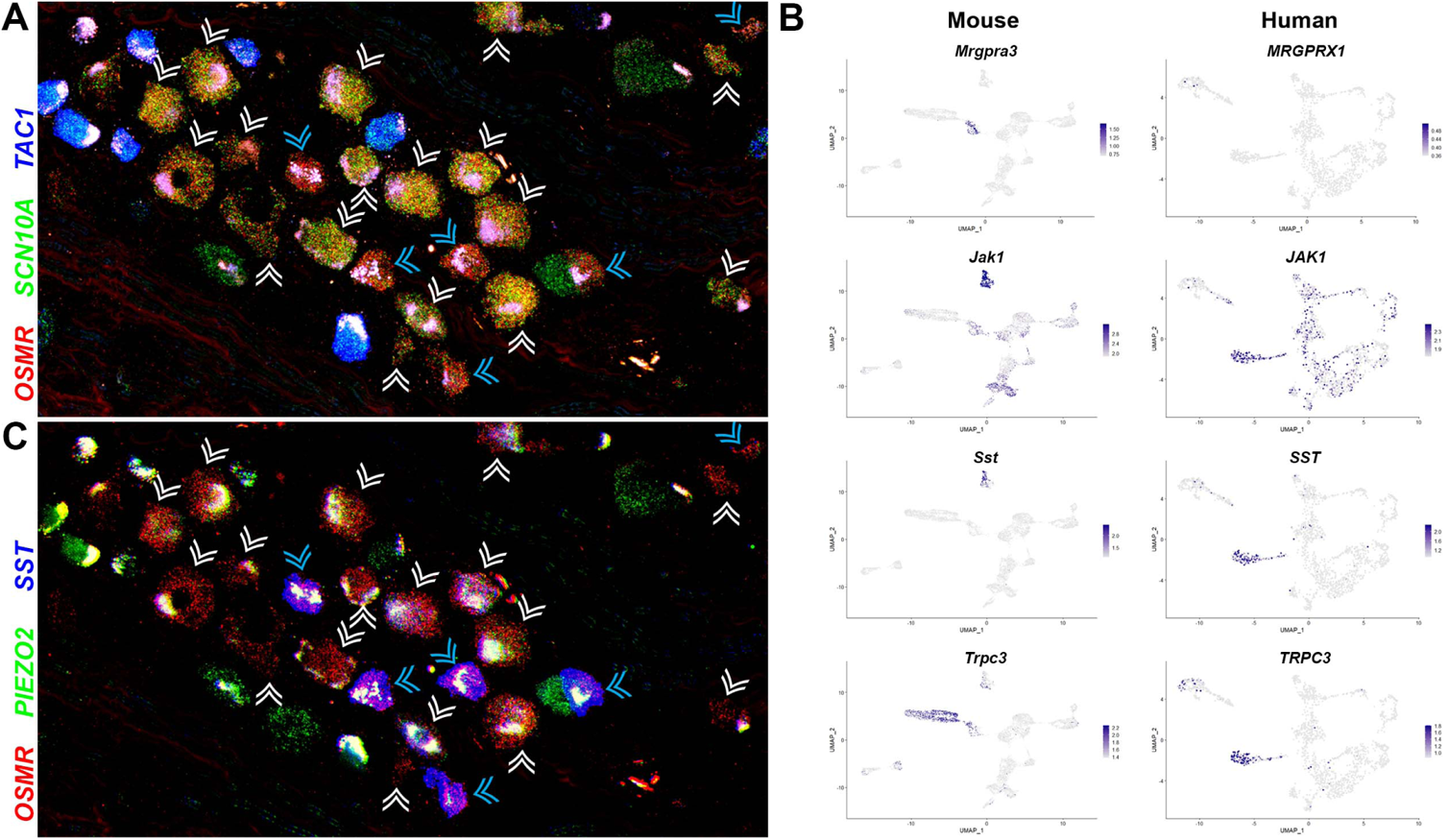
Two related classes of human non-peptidergic small diameter neurons that may mediate itch. (**A**) Confocal image of a section through a human DRG probed for nociception related genes using multiplexed ISH. In this view, many neurons expressing the itch related transcript, *OSMR* (red), are grouped together (arrowheads); cyan arrowheads point to cells that express relatively higher levels of *OSMR* than *SCN10A* (green). Peptidergic nociceptors marked by expression of *TAC1* (blue) and additional *SCN10A* positive cells are also present in this region of the ganglion. (**B**) UMAP representation of mouse and human DRG neurons showing relative expression level (blue) of genes that distinguish H10 and H11 and mark specific sets of mouse NP1-3 neurons. *MRGPRX1* is the human chloroquine receptor and the functional equivalent of *Mrgpra3*, which in mice marks NP2 cells. Note that co-expression patterns of *SST* and *JAK1* in H11 neurons resembles their expression in mouse NP3 pruriceptors but the ion channel *TRPC3* which also marks these cells is primarily expressed in mouse NP1 neurons; see Figure 5-figure supplement 1 for additional breakdown of similarities and differences between non-peptidergic neurons in mice and humans. (**C**) Confocal image of the group of candidate pruriceptors shown in (**A**) probed for expression of genes that distinguish Itch1 (*SST*, blue) from Itch2 (*PIEZO2*, green); note that neurons highlighted with cyan arrowheads have gene expression expected for Itch1 cells, whereas some Itch2 cells express lower levels of *SST* and also exhibit variation in the level of *PIEZO2* expression. Scale bars = 100 µm; see Figure 5-figure supplement 2 for individual channels and for expression of additional markers.

H10 cells are also distinguished from H11 and mouse pruriceptors by their prominent expression of the stretch gated ion channel *PIEZO2* (Figure 1C, Figure 2-figure supplement 1). The co-expression of itch related transcripts and this low threshold mechanosensor hint that H10 neurons may be responsible for the familiar human sensation known as mechanical itch. However, their relationship to NP1-neurons revealed by co-clustering mouse and human data (Figure 3) and their expression of markers for various other cell types (Figure 2-figure supplement 1, Figure 5-figure supplement 1) including non peptidergic cLTMRs suggest that their role in somatosensation may not be limited to itch alone.

A problem with single cell sequencing approaches is the sparse nature of the data making it difficult to disentangle expression level from proportional representation in any cluster. This means that except for the most highly expressed genes, there is inherent ambiguity in interpreting the expression patterns. ISH provides an independent and more analogue assessment of expression level that can help resolve this issue. Highly multiplexed ISH showed that *SST* divides the *OSMR* positive cells into two intermingled types (Figure 5C) in keeping with the sequence data (Figure 5B) and the relative expression patterns of *SCN10A* and *OSMR* (Figure 5A). Moreover, the prediction that *PIEZO2*-*OSMR* co-expression should mark *SST*-negative neurons was also largely borne out by ISH (Figure 5C, Figure 5-figure supplement 2). However, ISH also shows that some neurons expressing lower levels of *SST* are *PIEZO2*-positive and that some *OSMR*-positive H10 cells, contain only a very low level of the mechanosensory channel (Figure 5C, Figure 5-figure supplement 2). Therefore, H10 and H11 are by no means homogeneous populations and may not be as clearly distinguished from each other as snRNA sequencing suggests.

## Discussion

Comparison of single cell transcriptomic analysis of DRG neurons confirms that mouse and human somatosensory neurons express many of the same genes (*15*). However, although gross similarity in the transcriptomic classification of these cells can be discerned (peptidergic versus non-peptidergic; neurofilament rich, myelinated versus non-myelinated), the patterns of coordinated gene expression across species are not well conserved and both species exhibit unique specializations. Recently available transcriptomic data from the macaque (although likely biased to small diameter neurons) further highlights the individuality of somatosensory neurons across species (*40*). Surprisingly, in that study, despite major differences in gene expression between the two species, co-clustering approaches identified an apparently close relationship between cell types for both species (*40*). By contrast our analysis of human transcriptomic data using snRNA sequencing appears more quantitative in terms of neuron recovery but with lower read depth. Our transcriptomic data and analysis combined with highly multiplexed ISH provide strong evidence for major differences in small diameter nonpeptidergic neurons between humans and mice as well as the existence of other human specific cell-types. At one level, this interspecies variation was unexpected given that there is similarity between the neuronal types that comprise the mouse lumbar DRGs and trigeminal ganglia despite their very different types of innervation targets (*24, 25*). However, large changes in the receptive repertoire of other sensory systems have been observed and are thought to play a role in adaptation to specific ecological niches (*41*). Thus, the evolution of DRG receptor cell diversity further highlights the importance of appropriate sensory input for fitness and survival of a species. What is unusual relative to other senses is that transcriptomic differences are not limited to just the receptor repertoire for sensing environmental stimuli but instead extend to genes involved in the development and maintenance of defined neuronal subtypes. It is possible that this reflects major differences between mouse and human skin including fur covering. From a translational viewpoint, these differences could explain some of the problems in replicating results from mouse-based therapies (*22, 23*) in humans and the availability of the human data may help direct research towards new targets and even suggest precision medicine strategies (e.g. to treat cold pain).

The transcriptomic characterization of human somatosensory neurons presented here can also be compared with data that were recently obtained using spatial transcriptomics (*42*). The two types of analysis provide a very similar view of the classes of neurons present in human DRGs, strongly supporting the major differences between mouse and human somatosensory neurons. However, the sn-sequencing reveals some detail that goes beyond the spatial transcriptomic analysis. For example, the two populations of *TRPM8* expressing neurons that we describe and confirm using ISH were not distinguished in the spatial transcriptomic analysis (*42*). One reason for this difference may be the spatial grouping of transcriptomically similar neurons in human sensory ganglia (Figure 4) meaning that individual sections could present a biased view of neuronal types. Moreover, spatial transcriptomics does not directly sequence the individual neurons but instead targets areas of the section that often overlap neurons and surrounding tissue. Interestingly the spatial transcriptomic study highlighted sex differences in the transcriptome of individual clusters (*42*). Although we also examined male and female subjects, our data were not sufficiently powered to draw conclusions about sex differences since only a single male donor was studied. However, the major sex difference identified by spatial transcriptomics (*42*) revolved around the expression of *CALCA* in putative itch related cells. Tavares-Ferreira et al. (*42*) described a single itch cell class, resembling H10; they classified H11-like cells as silent nociceptors whereas our data imply a relationship with mouse NP3 neurons that have roles in pruriception (*7*). *CALCA* is expressed in H11 neurons (Figures 1, Figure 2-figure supplement 1). Therefore it will be necessary to carefully examine whether sex differences (*42*) correspond with gender related specialization, including perhaps a different ratio of H10 and H11 neurons or instead reflect the analytical method. Other markers for H10 and H11 including *GFRA2* and the ras-related estrogen-regulated growth inhibitor *RERGL*, which mark H10 cells (Figure 5-figure supplement 1), may help future efforts to resolve these issues. However, it should also be noted that our ISH analysis (Figure 5C) suggests that H10 and H11 cells are not homogeneous and exhibit some overlap in their expression of key genes.

Our analysis identified particularly surprising differences between small diameter non-peptidergic neurons in mice and humans. In mice, one distinctive subset of these cells are the cLTMRs that innervate hairy skin and are thought to be responsible for affective touch (*38*). At a transcriptomic level, humans do not have a clearly identifiable correlate for these cells although careful microneurography has revealed human c-fibers that respond to stroking (*43*). We suspect that some of these stroking responsive cells may be H10 neurons that unlike most mouse pruriceptors express high levels of *PIEZO2*, but it is also possible that some of these cells are other *PIEZO2*-expressing neurons that are also unique to humans (Figure 2B, Figure 1-figure supplement 3). Similarly, although H10 and H11 have some similarity to the mouse NP1-3 neurons, they also have major differences to all three types of cells. For example, in mice NP1 cells express a large combination of diagnostic markers (Figure 5-figure supplement 1) including *Mrgprd* that we did not find in our sequencing of human ganglion neurons. Bulk sequencing studies have identified *MRGPRD* expression in human DRG neurons (*44*), but recent ISH localization studies suggest broad but only low-level expression of this transcript together with *MRGPRX1* (*45*). This would fit with our co-clustering that identifies the *MRGPRX1*-expressing H10 neurons as related to NP1 cells. However, many of the other NP1 markers have potential roles in signal detection and transduction but are not H10 selective (Figure 5-figure supplement 1). Moreover, in mice, *Mrgpra3* (the functional equivalent of *MRGPRX1*) marks the distinct NP2 neurons.

Taken together, our results and analysis suggest that experiments in mice are likely to illustrate general principles that are important for sensory detection and perception in humans but also imply that specific details related both to genes and cell type responses may differ. In future studies, the central projections and targets of human somatosensory neuron subtypes might provide independent approaches for inferring function. Similarly, using immunohistochemistry to understand how these cell classes innervate the skin and other tissues may allow correlation of arborization patterns with microneurography results. Since microneurography can be complemented by microstimulation this could ultimately reveal the role of specific neuronal classes in sensory perception (*46*).

Our data provide a searchable database for gene expression in human DRG neurons. However, there are some limitations to the data and interpretation. For example, neither the number of neurons sequenced, nor the depth of sequencing is as comprehensive as for mice (*24, 25*). This means that rare neuronal subtypes and the expression patterns of moderately expressed genes may not be clear. Nonetheless, highly multiplexed ISH (Figures 2, 5) confirm the major findings both about cell-types and also gene expression and therefore substantiate the overall value of the data. The nuclear based sequencing approach used here has advantages in preventing gene expression changes during single cell isolation and is also likely to be less biased than cell-based approaches in terms of representation of the different cell-types (*2*). However, sn-RNA sequencing provides a somewhat distorted view of cellular gene expression, as has been described for sensory neurons in mice (*25*). Therefore, it will be important to confirm expression levels of specific genes using complementary approaches. Finally, any functional roles for neuronal classes identified here have been extrapolated from expression of markers and distant similarity to mouse counterparts. Given the extensive differences that we report, some of these conclusions may need to be revised once cell class can be linked to neuronal function in human subjects.

## Materials and Methods

### Study design

Transcriptomic analysis of human DRG neurons was carried out to establish similarities and differences between human somatosensory neurons and their counterparts in model organisms and to provide a resource. We chose a nuclear based strategy because of its simplicity and quantitative nature relative to isolation of cells (*25*). All tissue was obtained from deidentified organ donors and was not pre-selected or otherwise restricted according to health conditions. We used DRGs from both male and female donors for the sequencing and ISH localization experiments. Randomization and blinding were not used because of the nature of our experiments. Similarly, before starting this study, we had no relevant information for setting sample size for snRNA sequencing from human DRG neurons. Therefore, we stopped data collection when we empirically determined that the cost of adding extra data outweighed the benefit of additional sequencing. In essence, numbers of sn-transcriptomes analyzed were limited by the availability of material and the difficulty of isolating human DRG nuclei with preservation of their transcriptome. We considered that the dataset would serve as a valuable and relatively comprehensive resource once including additional material from an individual preparation made only minor differences to the pattern of clustering we observed. Criteria for data exclusion followed standards in the field (see below) and sample sizes and numbers of replicates are also typical for this type of study and are described in the relevant experimental sections. Apart from the exclusions described for single nucleus experiments, all data obtained were included in our study.

### Isolation of human DRG nuclei

DRG-recovery was reviewed by the University of Cincinnati IRB #00003152; Study ID: 2015-5302, title Human dorsal root ganglia and was exempted. Lumbar L4 and L5 DRGs were recovered from donors withing 90 minutes of cross-clamp (*47*). For RNA sequencing, human DRGs immediately were cut into 1-2 mm pieces and stored in RNA-later (ThermoFisher, Cat# AM7021). For ISH, DRGs were immersion fixed in 4% paraformaldehyde in phosphate buffered saline (PBS) overnight and cryoprotected in 30% sucrose and were frozen in OCT (Tissue Tek). Excess RNA-later was removed and the tissues were frozen on dry ice and stored at −80 °C. Nuclei were isolated from each donor separately as described previously (*25*) with minor modification. Briefly, the tissues were homogenized with a Spectrum Bessman tissue pulverizer (Fisher Scientific, CAT# 08-418-3) in liquid nitrogen. The sample was then transferred to a Dounce homogenizer (Fisher Scientific, Cat# 357538) in 1 ml of freshly prepared ice-cold homogenization buffer (250 mM sucrose, 25 mM KCl, 5 mM MgCl_2_, 10 mM Tris, pH 8.0, 1 µM DTT, 0.1% Triton X-100 (v/v). To lyse cells and preserve nuclei homogenization used five strokes with the ‘loose’ pestle (A) and 15 strokes with the ‘tight’ pestle (B). The homogenate was filtered through a 40 µm cell strainer (ThermoFisher, cat# 08-771-1), was transferred to low bind microfuge tubes (Sorenson BioScience, cat# 11700) and centrifuged at 800 g for 8 mins at 4°C. The supernatant was removed, the pellet gently resuspended in 1 ml of PBS with 1% BSA and SUPERaseIn RNase Inhibitor (0.2 U/µl; ThermoFisher, Cat#AM2696) and incubated on ice for 10 min.

Neuronal nuclei selection was performed by incubating the sample with a rabbit polyclonal anti-NeuN antibody (Millipore, cat#ABN78) at 1:4000 with rotation at 4°C for 30 min. The sample was then washed with 1 ml of PBS with 1% BSA and SUPERaseIn Rnase Inhibitor and centrifuged at 800 g for 8 mins at 4°C. The resulting pellet was resuspended in 80 µl of PBS, 0.5% BSA, 2 mM EDTA. 20 µl of anti-rabbit IgG microbeads (Miltenyi biotec, cat# 130-048-602) were added to the sample followed by a 20 min incubation at 4°C. Nuclei with attached microbeads were isolated using an LS column (Miltenyi Biotec, cat# 130-042-401) according to the manufacturer’s instruction. The neuronal nuclei enriched eluate was centrifuged at 500 g for 10 min, 4°C. The supernatant was discarded, and the pellet was resuspended in 1.5 ml of PBS with 1% BSA. To disrupt clumped nuclei, the sample was homogenized on ice with an Ultra-Turrax homogenizer (setting 1) for 30 secs. An aliquot was then stained with trypan blue and the nuclei were counted using a hemocytometer. The nuclei were pelleted at 800 g, 8 mins at 4°C and resuspended in an appropriate volume for 10X Chromium capture. A second count was performed to confirm nuclei concentration and for visual inspection of nuclei quality.

### Single nuclear capture, sequencing, and data analysis

10X Chromium capture and library generation were performed according to manufacturer’s instructions using v3 chemistry kits. Next generation sequencing was performed using Illumina sequencers. 10X Chromium data were mapped using CellRanger to a pre-mRNA modified human genome (GRCh38.v25.premRNA). Data analysis used the Seurat V3 packages developed by the Satija lab and followed standard procedures for co-clustering (*28*). For sn-RNA sequencing experiments cell filtering was performed as follows: outliers were identified and removed based on number of expressed genes and mitochondrial proportion as is standard practice in single cell transcriptomic analysis. After initial co-clustering of data from the different preparations, non-neuronal cell clusters were identified by their gene expression profiles: clusters not expressing high levels of neuronal or somatosensory genes like *SNAP25*, *SCN9A*, *SCN10A*, *PIEZO2*, *NEFH* etc. but instead expressing elevated levels of markers of non-neuronal cells including *PRP1*, *MBP* and *APOE* were tagged as non-neuronal and were removed to allow re-clustering of “purified” human DRG neurons. A total of 1837 human DRG neuronal nuclei were included in the analysis. The mean number of genes detected per nucleus was 2839 (range 501 – 9652), with a standard deviation of 1917. Moderate changes in clustering parameters and in the cutoffs for data inclusion/exclusion as well as leaving out nuclei from any single preparation made differences in how the data were represented graphically but not to the main conclusions. All the different transcriptomically related neuron-types described here could still be readily discerned in UMAP analysis of expression data.

For analysis of the mouse, a random subset of data from sn-RNA sequencing of DRGs from wild type mice was extracted from data deposited by the Woolf lab (*29*). The data were filtered according to gene count and mitochondrial DNA leaving approx. 7500 cells that were clustered using standard methods (*28*). The expression patterns that are described for genes in mice can also be checked in the outstanding and easy to search single cell analysis provided by Sharma et al. (*24*). Co-clustering of mouse and human data used methods described by the Satija lab (*28*). We experimented using different numbers of mouse neurons but found broadly similar results over a range from 1,800 to 7,500 mouse neurons. At lower numbers of mouse neurons, the same relationships as shown in Figure 3 could be discerned but both the mouse and human neurons were less well organized. When we used substantially greater numbers of mouse neurons from the full Renthal dataset (*29*) only co-clustering of large diameter neurons across species was observed. 30 principal components best describing the data were calculated based on the integrated mouse and human expression data and used as the basis for clustering and UMAP projection. In order to quantify the similarity/dissimilarity between a given mouse cluster and any human cluster, the Kullback-Leibler divergence between their distributions in 30-dimensional continuous space was estimated as described previously (*48*) using R (Code available here). The natural logarithm of the Kullback-Leibler divergences for each mouse/human pairing was plotted as a heatmap in R.

### In situ hybridization

Cryosections from human DRGs were cut at 20 µm and used for ISH with the RNAscope HiPlex Assay (Advanced Cell Diagnostics) following the manufacturer’s instructions. The following probes were used: *NEFH* (cat# 448141); *TRPM8* (cat# 543121); *PIEZO2* (cat# 449951); *SCN10A* (cat# 406291); *NTRK2* (cat# 402621); *TAC1* (cat# 310711); *OSMR* (cat# 537121); *SST* (cat# 310591); *TRPV1* (cat# 415381).

Confocal microscopy (5 µm optical sections) was performed with a Nikon C2 Eclipse Ti (Nikon) at 40X magnification. All confocal images shown are collapsed (maximum projection) stacks. HiPlex images were aligned and adjusted for brightness and contrast in ImageJ as previously described (*2*). Diagnostic probe combinations were used on at least three sections from at least two different individuals with qualitatively similar results. Overall, we used sections from ganglia from 4 different donors, however, the signal intensity of all probes varied between the individual ganglia making identification of positive signal risky in some cases. Strongest signals were observed for sections from an 18-year-old male and a 35-year-old female donor. All images displayed here, and our analysis including cell counts were from sections of ganglia isolated from these two individuals. When individual channels are displayed in the supplements, the strongest autofluorescence signals have been selected and superimposed using photoshop to help focus attention on ISH-signal.

### Spatial analysis of cell clusters

In order to quantify spatial clustering of cell types, neurons in ISH images (1 male, 1 female) were manually outlined and annotated as either *NEFH*-only, *SCN10A*-only or as expressing both. Centroid coordinates of these cells and their distances were analyzed in Python 3.7. For each of the 346 *NEFH* cells and *244 SCN10A* cells, the nearest neighbors were identified based on Euclidean distance (Scikit-Learn package) and the percentage of *NEFH* and *SCN10A* cells in each neighborhood of size 1 - 40 cells was calculated. Statistical significance between *NEFH*-surrounding and *SCN10A*-surrounding neighborhoods was determined using a one-tailed Mann-Whitney U test (Scipy Stats package).

## Acknowledgments

We thank the Genomics and Computational Biology Core (National Institute on Deafness and Other Communication Disorders) for sequencing; this work utilized the computational resources of the NIH HPC Biowulf cluster (http://hpc.nih.gov). We also thank LifeCenter, Cincinnati and the donor families for their generosity. We are also indebted to Drs. Mark Hoon, Claire Le Pichon and Alexander Chesler and members of our groups for valuable suggestions

## Author contributions

Conceptualization: MQN, NJPR, SD; Methodology: MQN, LJvB, ANR, NJPR, SD; Data curation: MQN, NJPR; Data analysis: MQN, LJvB, NJPR; Writing-original draft: NJPR; Writing-review and editing: MQN, LJvB, ANR, NJPR, SD.

## Data and materials availability

Sequence data will be available on publication in GEO, accession number GSE168243; a searchable version of the data will also be made available at a permanent address; currently gene-expression data can be searched at: https://lars-von-buchholtz.shinyapps.io/shinyseurat/

## Supplementary Figures

**Figure 1-figure supplement 1.**
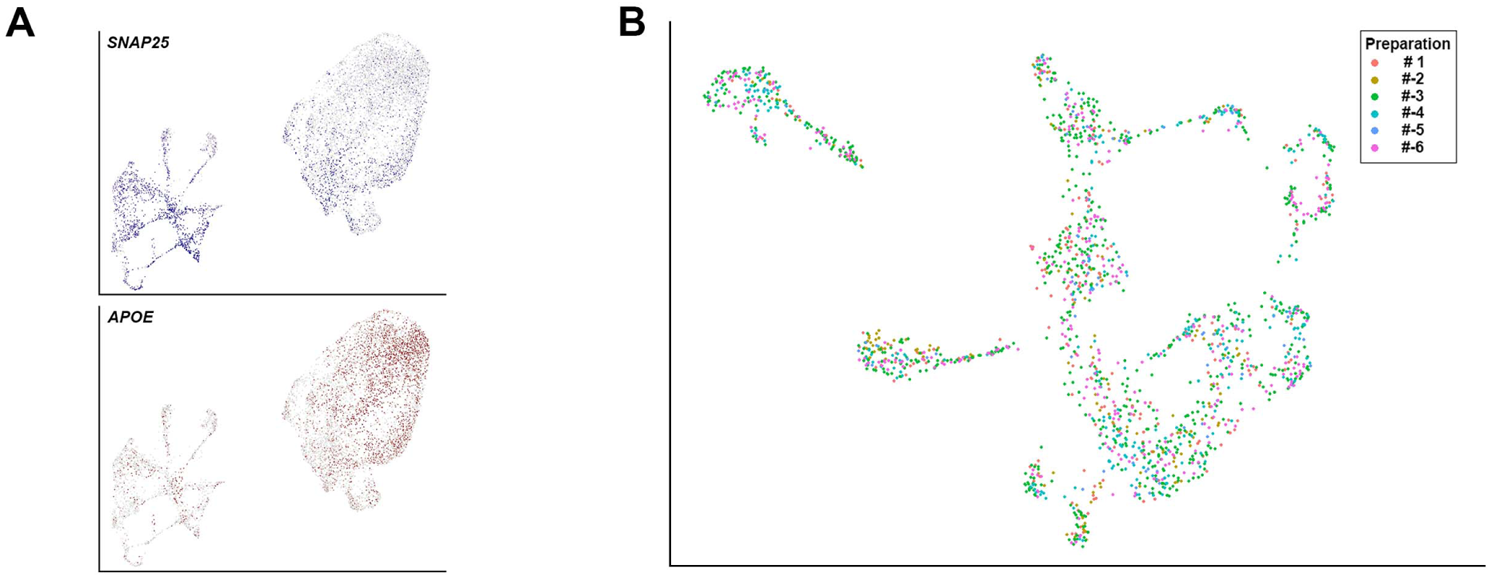
Support for the clustering of human DRG neurons. (**A**) UMAP representation showing relative expression levels of the neuronal marker *SNAP25* (blue) and the non-neuronal gene *APOE* (red) in the initial clustering of sn-RNA sequencing data. It should be noted that the majority of the non-neuronal cells came from a single nuclear isolation where it is likely that neuronal nuclear purification was not effective. Sequence analysis indicated that most of the non-neuronal cells were satellite microglia although other cell types were also seen. (**B**) UMAP representation of the clustering of human DRG neurons highlighting the contributions of the six different preparations to the dataset. Note that clusters were populated with data from multiple different preparations.

**Figure 1-figure supplement 2.**
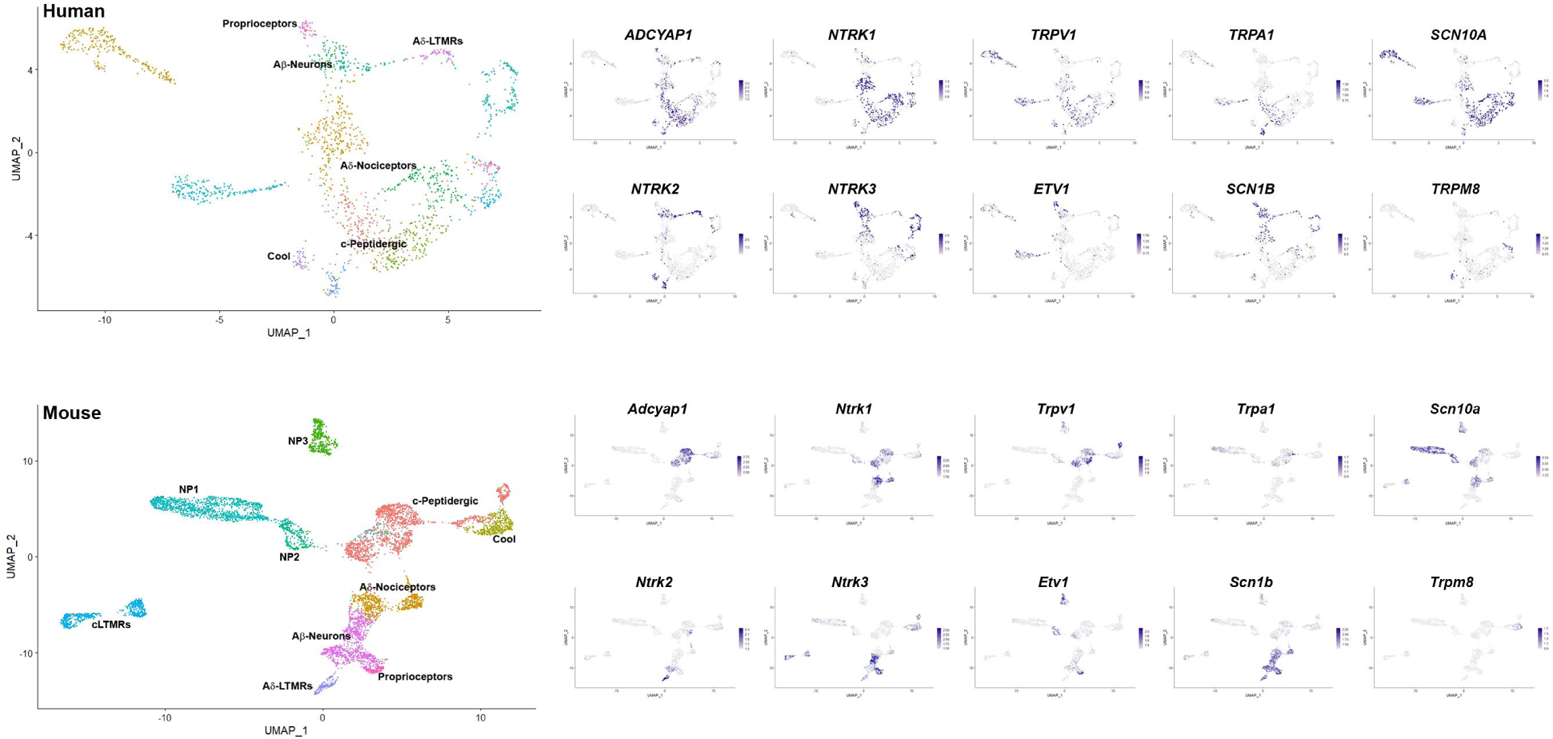
Additional markers that support similarities between DRG neuronal clusters across species. UMAP representations: upper panels, human sn-RNA sequencing; lower panels mouse sn-RNA sequencing. To the left, identity of the different neuronal types is differentially colored. For the mouse, the identities of all clusters are indicated, for the human data, the position of names indicate the clusters with shared markers indicating a match to mouse neuronal types. The expression pattern of several such marker genes (blue) is shown to the right (see also Figure 1 and Figure 1-figure supplement 3).

**Figure 1-figure supplement 3.**
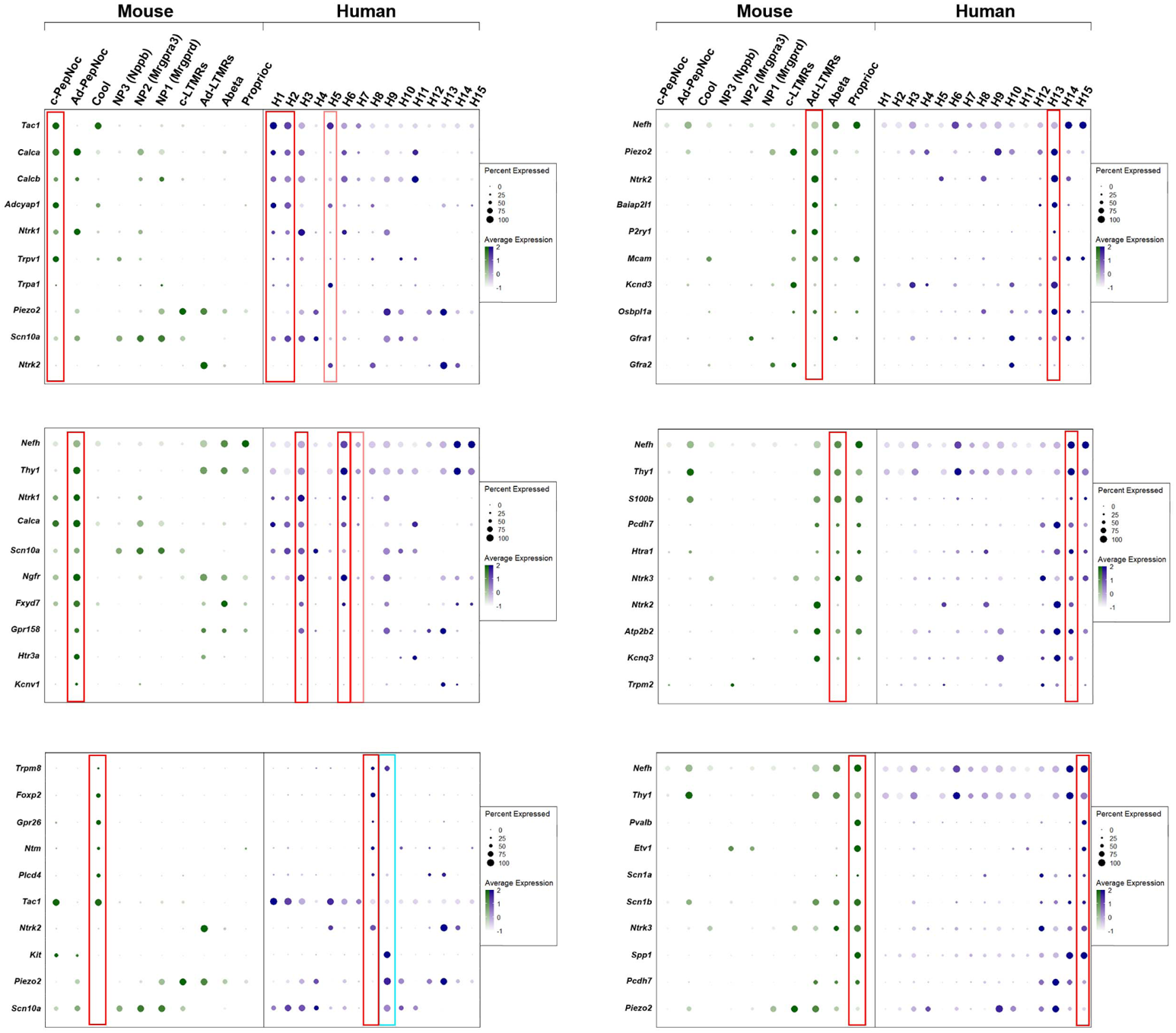
Dotplots of gene expression supporting similarity of several DRG neuronal classes in mice and humans. Dotplots displaying information about the fractional expression and relative expression level of marker genes in the different identity classes of mouse (left, gray-green scale) and human (right, gray-blue scale). Marker genes were chosen based on their expression (or lack of expression) in particular classes of mouse neurons. Red boxes highlight the relationship between a particular mouse class and human neurons; fainter red boxes indicate human classes that share some characteristics with the highlighted mouse neurons. Left panel top to bottom: c-peptidergic nociceptors (c-PepNoc); Aδ-peptidergic nociceptors (Ad-PepNoc); cool sensing cells (Cool). Right panel top to bottom: Aδ-LTMRs (Ad-LTMRs); Aβ-neurons (Abeta); proprioceptors (Proprioc). The cyan box highlights H9 a human specific cell type that expresses *TRPM8* but also nociceptor markers and the mechanosensitive channel *PIEZO2* unlike the putative cool sensing cells in human (H8) and in mice.

**Figure 2-figure supplement 1.**
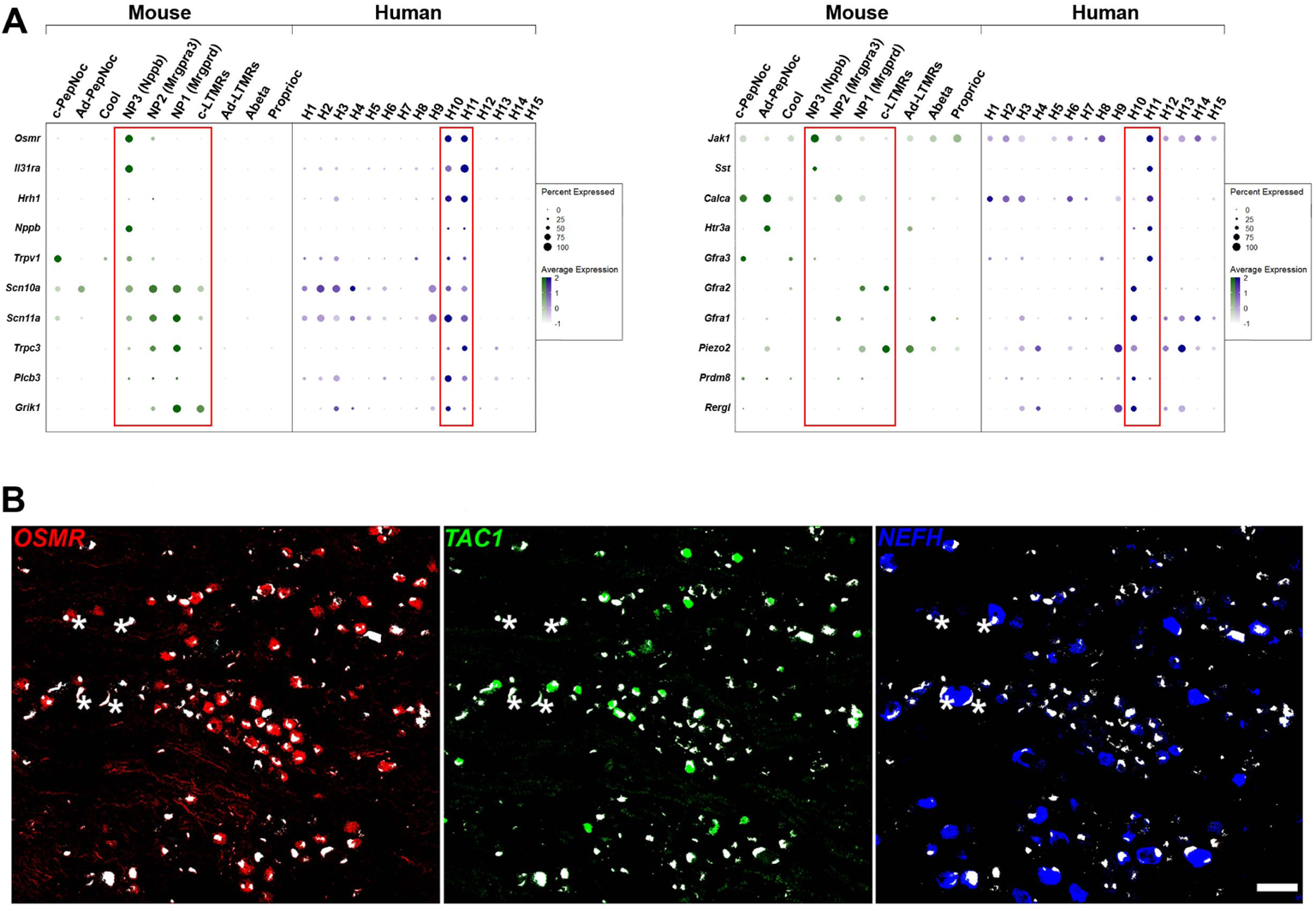
H10 and H11 are classes of human DRG neurons that express a range of itch related genes. **(A)** Dotplots displaying information about the fractional expression and relative expression level of marker genes in the different identity classes of mouse (left, gray-green scale) and human (right, gray-blue scale). Left panel, a selection of genes with relatively similar expression in H10 and H11 cells includes functionally relevant genes that tune mouse NP3 cells to respond to itch. Right panel shows genes that distinguish H10 and H11; some of these are also markers for mouse NP3 cells, others are more highly expressed in the other classes of mouse small diameter non-peptidergic neurons. Red boxes highlight H10 and H11 and the four classes of mouse small diameter non-peptidergic neurons. **(B)** Individual channels for the ISH shown in Figure 2D highlight *NEFH* as a marker for large diameter neurons and further emphasize the relatively limited overlap between these three probes. Strong autofluorescence signals that are present in all channels have been masked in white; four examples of autofluorescence (where there is also signal for one probe in that cell) are highlighted by stars.

**Figure 4-figure supplement 1.**
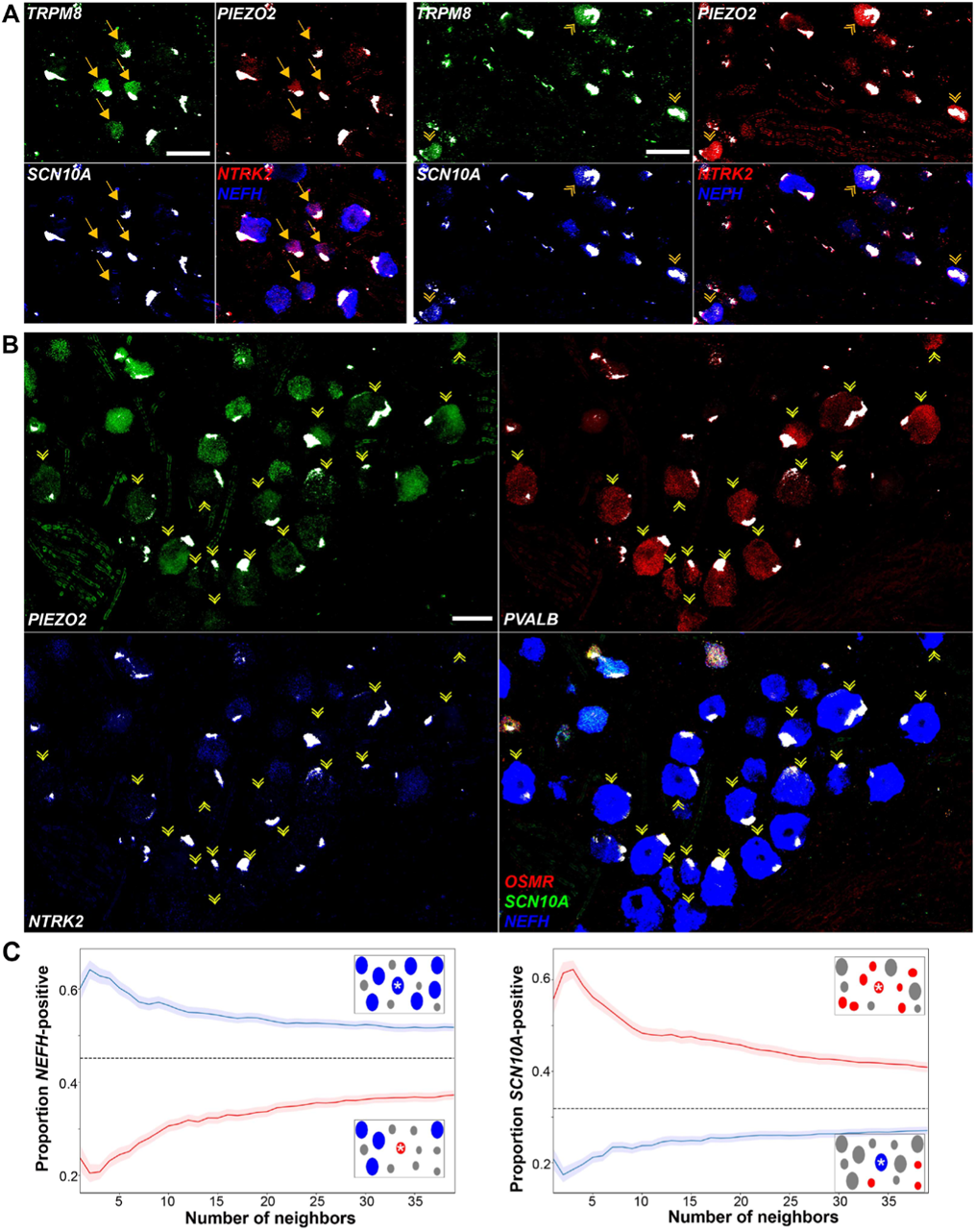
Expression profiles of clusters of H8 (cool), H9 (human specific) and H15 (proprioceptive) neurons. Individual channels for the ISH images in Figure 4 are shown to highlight the expression patterns described in the text. Strong autofluorescence signals that are present in all channels have been masked in white. Additional combinations of gene expression data in these regions of the ganglion are shown to identify neurons and reveal extra transcriptomic information. (**A**) Spatial clusters containing H8 putative cool sensing neurons (left) and H9 cool/mechanosensory nociceptors (right) are shown and identified as in Figure 4A. H8 neurons are generally *NTRK2* positive but are only weakly positive for *NEFH*. By contrast, H9 cells are *NEFH* positive but do not express significant *NTRK2*, in keeping with transcriptomic data. Note that clusters of these neurons segregate in distinct fields of the ganglion. (**B**) H15 (presumptive proprioceptors) form a sub-cluster of *NEFH*-positive cells that are essentially devoid of nociceptors (marked by *SCN10A*) including non-peptidergic neurons expressing *OSMR*. Many of the *PVALB*-negative large diameter neurons in this region of the ganglion were *NTRK2* positive; by contrast this gene was not detectable in H15 cells in keeping with the transcriptomic data. Scale bars = 100 µm; arrows and arrowheads are as in Figure 4. **(C)** All labeled neurons in Figure 4C were identified and were scored as *NEFH*, *SCN10A* or double positive. The nearest n neighbors were identified (for n = 1 - 40): blue lines represent proportion (mean, solid line ± s.e.m., shaded) of cells surrounding *NEFH*-only neurons; red lines, proportion (mean, solid line ± s.e.m., shaded) of cells surrounding *SCN10A*-only neurons. Left panel: proportion of surrounding cells that were only *NEFH*-positive. Right panel: proportion of surrounding cells that were only *SCN10A*-positive. Dashed black lines are the proportions expected for randomly distributed cells. Insets schematically show a central cell (highlighted by a star) and the surrounding neurons. Neighboring *NEFH*-only cells are colored blue and *SCN10A*-only cells are colored red when these are being scored in the associated graph; grey cells are either single positive for the other marker or double positive neurons. Clustering was statistically significant across the complete range (1 – 40 neighbors) maximum p < 1.3 x 10^-18^ (one-tailed Mann-Whitney U-test); n = 244 *SCN10A*-only and 346 *NEFH*-only cells confirming both short and long-range grouping of similar classes of human DRG neurons (sections from 2 donors were analyzed).

**Figure 5-figure supplement 1.**
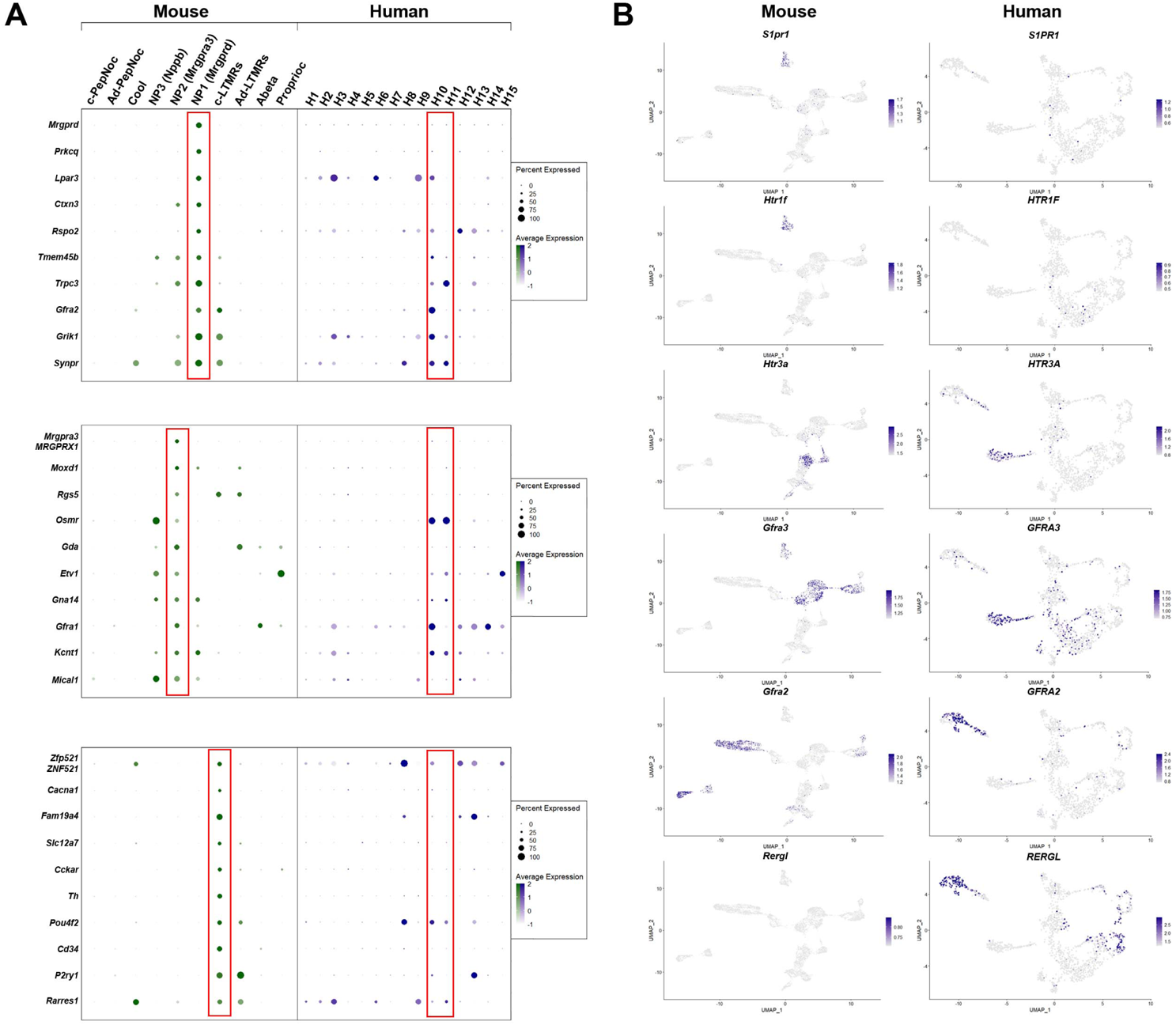
Gene expression patterns in H10 and H11 classes of human DRG neurons are distinct from classes of mouse small diameter non peptidergic neurons. **(A)** Dotplots displaying information about the fractional expression and relative expression level of marker genes in the different identity classes of mouse (left, gray-green scale) and human (right, gray-blue scale). From top to bottom, markers of NP1, NP2 and c-LTMRs in mice generally show only limited expression in the presumptive human non-peptidergic nociceptors H10 and H11. NP1 selective genes confirm greater similarity of H10 neurons to this type of mechanonociceptor. Right panels UMAP representation of mouse and human DRG neurons showing relative expression level (blue) of several genes that highlight differences between H10 and H11 neurons and potential mouse counterparts. **(B)** UMAP representation of mouse and human DRG neurons showing relative expression level (blue) of several genes highlighting differences between H10 and H11 neurons and potential mouse counterparts.

**Figure 5-figure supplement 2.**
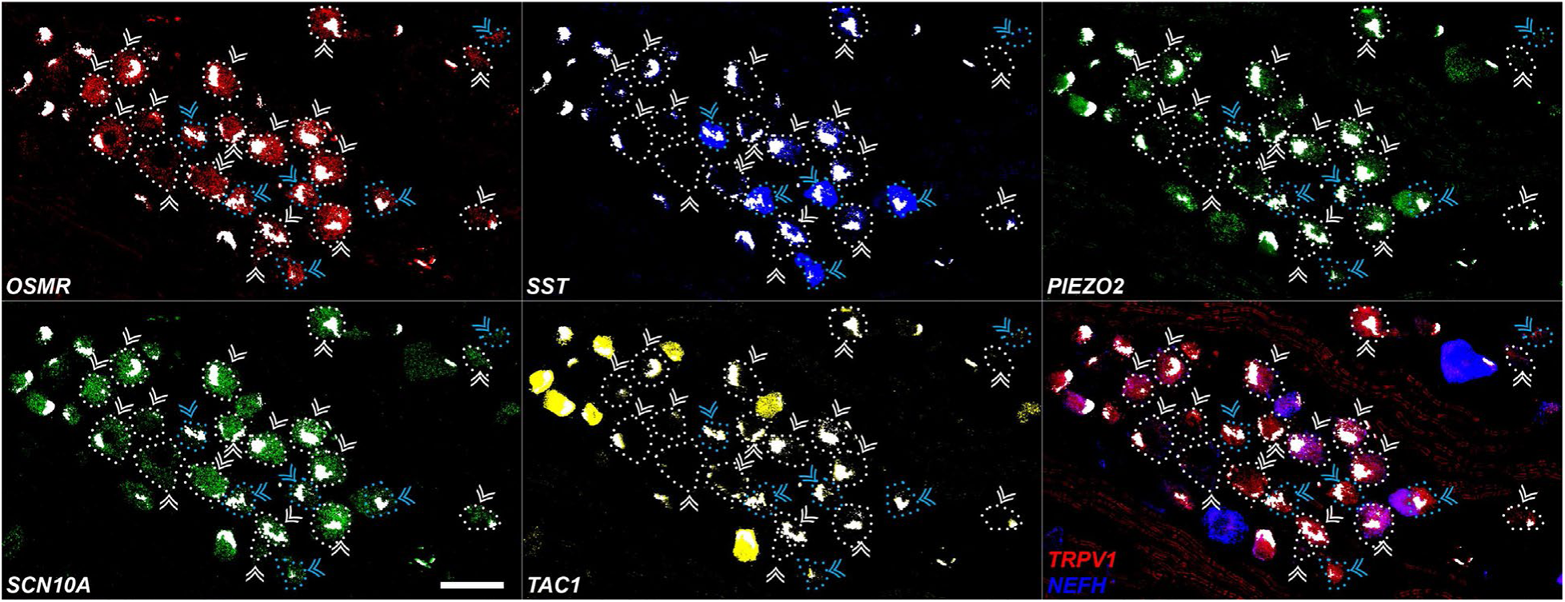
Expression profiles of human H10 and H11 DRG neurons. Individual channels for the ISH images in Figure 5 are shown to highlight the expression patterns described in the text. Strong autofluorescence signals that are present in all channels have been masked in white. In addition, ISH reveals that H10 and H11 neurons also express *TRPV1* but as expected from the transcriptomic data at most express very low levels of *NEFH*. Scale bar = 100 µm; arrowheads are as in Figure 5; to further highlight the relevant cells, *OSMR*-positive cells are highlighted by dotted outlines that match the coloring of arrowheads.

